# Molecular Mechanism Underlying Inhibition of Intrinsic ATPase Activity in a Ski2-like RNA Helicase

**DOI:** 10.1101/758169

**Authors:** Eva Absmeier, Karine F. Santos, Markus C. Wahl

**Author notes:** These authors contributed equally to this work (Eva Absmeier, Karine F. Santos). MRC Laboratory of Molecular Biology, Cambridge Biomedical Campus, Cambridge CB2 0QH, UK. Sphingotec GmbH, Neuendorfstr. 15A, 16761 Henningsdorf, Germany (Karine F. Santos).

## Abstract

RNA-dependent NTPases can act as RNA/RNA-protein remodeling enzymes and typically exhibit low NTPase activity in the absence of RNA/RNA-protein substrates. How futile intrinsic NTP hydrolysis is prevented is frequently not known. The ATPase/RNA helicase Brr2 belongs to the Ski2-like family of nucleic acid-dependent NTPases and is an integral component of the spliceosome. Comprehensive nucleotide binding and hydrolysis studies are not available for a member of the Ski2-like family. We present crystal structures of *Chaetomium thermophilum* Brr2 in the apo, ADP-bound and ATPyS-bound states, revealing nucleotide-induced conformational changes and a hitherto unknown ATPyS binding mode. Our results in conjunction with Brr2 structures in other molecular contexts reveal multiple molecular mechanisms that contribute to the inhibition of intrinsic ATPase activity, including an N-terminal region that restrains the RecA-like domains in an open conformation and exclusion of an attacking water molecule, and suggest how RNA substrate binding can lead to ATPase stimulation.

**HIGHLIGHTS:**

- Crystal structures of Brr2 in complex with different adenine nucleotides.
- The Brr2 N-terminal region counteracts conformational changes induced by ATP binding.
- Brr2 excludes an attacking water molecule in the absence of substrate RNA.
- Different helicase families resort to different NTPase mechanisms.

## INTRODUCTION

RNA-dependent NTPases represent a large group of enzymes that are involved in diverse aspects of gene expression and gene regulation in all domains of life. Most RNA-dependent NTPases belong to superfamilies (SFs) 1 or 2 of nucleic acid-dependent NTPases (Jankowsky, 2011). SF1 and SF2 enzymes comprise a core of dual RecA-like motor domains that can bind and hydrolyze NTPs and that can bind RNA or RNA-protein (RNP) substrates dependent on the NTP-bound state (Jankowsky and Fairman, 2007). NTP binding, hydrolysis, product release and rebinding elicit conformational changes, with different conformational states exhibiting different RNA/RNP affinities. As a consequence, many of these enzymes can bind, deform and release RNAs/RNPs or translocate on RNAs in an NTPase-dependent manner to achieve, e.g., RNA duplex unwinding or disruption of RNPs (Singleton et al., 2007).

The core RecA-like domains of SF1 and SF2 enzymes contain up to twelve conserved sequence motifs that mediate and functionally couple the NTP/RNA/RNP transactions (Fairman-Williams et al., 2010). In addition, many SF1 and SF2 enzymes contain additional domains inserted into or appended to the core domains, forming helicase units (cassettes), in which the helicase-associated domains facilitate NTP/RNA/RNP-related activities of the dual-RecA cores and modulate the precise molecular mechanisms, by which RNA/RNP remodeling is achieved (Jankowsky, 2011). Based on which motifs are present, the exact sequences of the motifs and the presence of additional domains, SF1 and SF2 members are divided into several families each (Fairman-Williams et al., 2010). For instance, SF2 comprises ten families, of which five (Ski2-like, RIG-I-like, DEAD-box, DEAH/RHA and NS3/NPH-II) contain RNA-dependent NTPases (Fairman-Williams et al., 2010).

To prevent futile NTP hydrolysis, the NTPase activities of RNA-dependent NTPases are auto-inhibited in the absence of substrate RNAs/RNPs. However, for many SF1 and SF2 enzymes the precise mechanisms, by which intrinsic NTPase activity is held at check, are presently not known. Elucidation of such mechanisms requires the determination of atomic-level 3D structures of the enzymes in different NTP and RNA/RNP substrate-bound states. Detailed NTP binding studies have been conducted for members of the DEAD-box (Putnam and Jankowsky, 2013), DEAH/RHA (Tauchert et al., 2017) and NS3/NPH-II (Gu and Rice, 2010) families of SF2 enzymes. However, a similarly comprehensive analysis of a Ski2-like family member has so far not been reported.

The ATP-dependent RNA helicase Brr2 is a member of a small subgroup of Ski2-like enzymes that comprise a tandem repeat of Ski2-like helicase cassettes (Figure 1A). Brr2 is an integral component of the spliceosome and is required for the remodeling of an initially assembled, pre-catalytic spliceosome (B complex) into a catalytically competent spliceosome (Xu et al., 1996; Zhang et al., 2015), during which Brr2 disrupts the U4/U6 di-small nuclear (sn) RNP by unwinding the U4/U6 di-snRNAs (Bertram et al., 2017; Laggerbauer et al., 1998; Nguyen et al., 2016; Plaschka et al., 2018; Raghunathan and Guthrie, 1998; Zhan et al., 2018). As in all Ski2-like enzymes, the two RecA-like domains of the Brr2 helicase cassettes are followed by a winged-helix (WH) domain, a helical bundle (HB) or “ratchet” domain and a helix-loop-helix (HLH) domain; the Brr2 helicase cassettes additionally comprise a C-terminal immunoglobulin-like (IG) domain (Figure 1A). While both Brr2 cassettes can bind adenine nucleotides (Santos et al., 2012), only the N-terminal cassette (NC) can hydrolyze ATP and couple ATP hydrolysis to RNA duplex unwinding or RNP disruption; the C-terminal cassette (CC) is inactive as an ATPase/helicase (Kim and Rossi, 1999; Santos et al., 2012).

**Figure 1.**
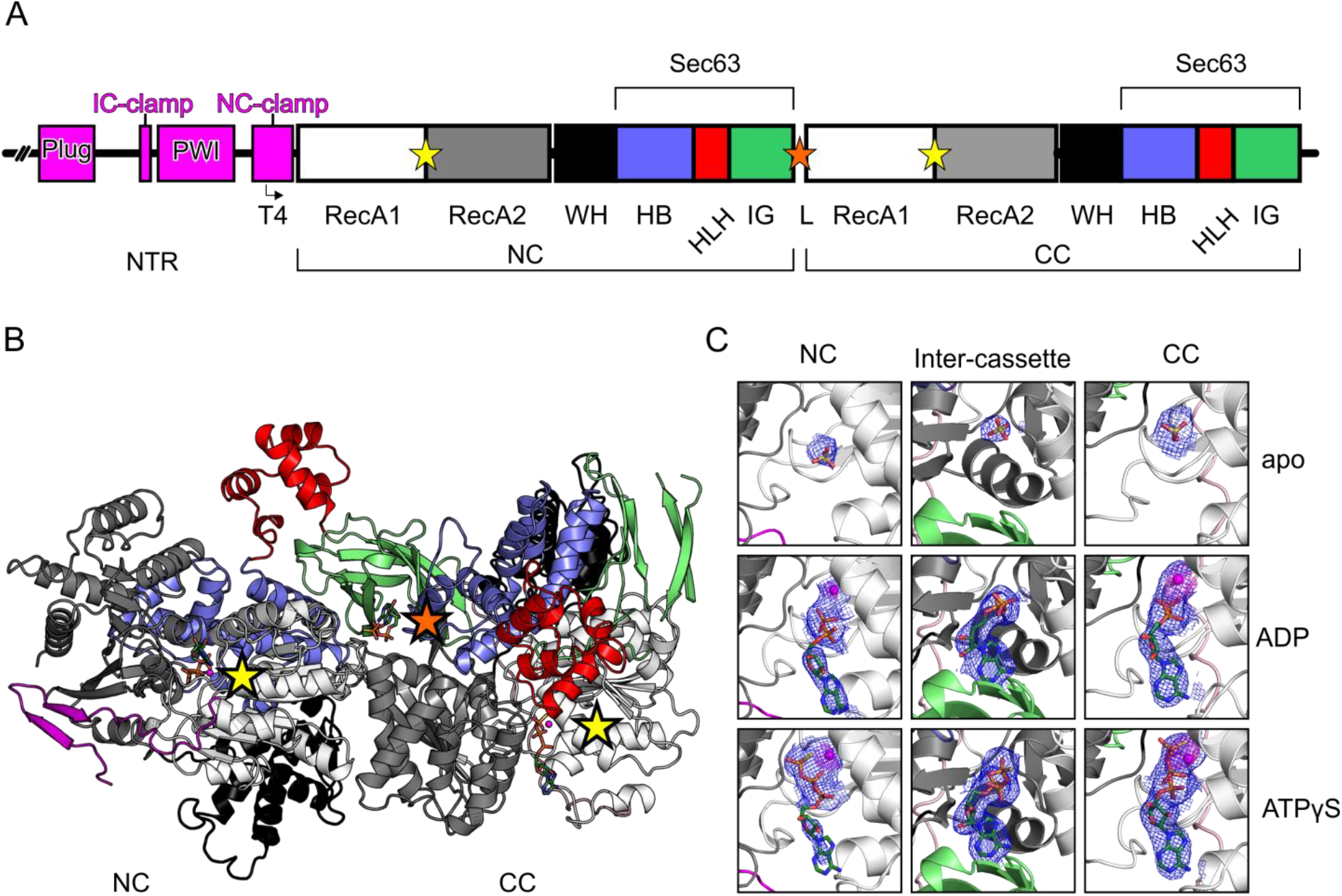
Brr2 Domain Organization and Nucleotide Binding. (A) Scheme showing the Brr2 domain organization. The angled arrow indicates the starting position of cBrr2^T4^. NTR, N-terminal region; plug, plug domain; PWI, PWI-like domain, IC-clamp, inter-cassette clamp; NC-clamp, N-terminal cassette clamp; NC, N-terminal cassette; CC, C-terminal cassette; WH, winged helix domain; HB, helical bundle domain; HLH, helix-loop-helix domain; IG, immunoglobulin-like domain; L, linker; Sec63, Sec63 homology unit. Yellow stars indicate the canonical nucleotide binding pockets between the RecA domains in the NC and CC. The orange star indicates the position of the third nucleotide bound between the NC and CC. (B) Overall structure of cBrr2^T4^ bound to ATPγS. Domains/regions are colored as in (A). (C) Composite 2F_o_-F_c_ omit maps (blue; contoured at the 1 σ-level) of the bound sulfate ions and nucleotides, and anomalous difference Fourier maps (magenta; contoured at the 4 σ-level) showing positions of Mn^2+^ ions (magenta spheres). Sulfate ions, ADP and ATPγS are shown as sticks and colored by atom type (carbon, dark green; nitrogen, blue; oxygen, red; phosphorus, orange; sulfur, yellow).

The two helicase cassettes in Brr2 are preceded by a ~450-residue N-terminal region (NTR), that encompasses two folded domains (“plug” and PWI-like) and adjacent unstructured regions (Absmeier et al., 2015b, 2015a) (Figure 1A). Functional analyses of stepwise N-terminally truncated *Chaetomium thermophilum* Brr2 (cBrr2) have shown that the NTR has an inhibitory effect on Brr2 ATPase, RNA-binding and RNA-unwinding activities *in vitro* (Absmeier et al., 2015b). *In vivo*, the NTR is essential for yeast viability, stable association of Brr2 with the U4/U6-U5 tri-snRNP and tri-snRNP stability (Absmeier et al., 2015a; Zhang et al., 2015). Recent cryo-electron microscopy (cryoEM) structures of yeast U4/U6•U5 tri-snRNPs (Agafonov et al., 2016; Nguyen et al., 2016; Wan et al., 2016) and of pre-catalytic B complex spliceosomes (Bertram et al., 2017; Plaschka et al., 2018; Zhan et al., 2018) showed that the NTR is completely detached from the helicase cassettes to allow Brr2 to engage the U4 snRNA strand of the U4/U6 di-snRNP.

Here, we describe crystal structures of a cBrr2 N-terminal truncation variant in the apo, ADP-bound and ATPyS-bound states. Comparison with members of the other SF2 helicase families revealed an unusual pre-catalysis conformation in Brr2. We also compared our structures to the Brr2 subunits in cryoEM structures of the yeast tri-snRNP (Nguyen et al., 2016) and pre-catalytic spliceosomal B complexes (Bertram et al., 2017; Plaschka et al., 2018; Zhan et al., 2018), providing a molecular explanation for Brr2’s low intrinsic ATPase activity and for how ATPase activity is stimulated once the enzyme engages a RNA/RNP substrate.

## RESULTS

### Structures of Nucleotide-Bound cBrr2^T4^

Removal of almost the entire NTR (residues 1-472) led to a cBrr2 variant (cBrr2^T4^; Figure 1A) with enhanced intrinsic and RNA-stimulated ATPase activities compared to full-length cBrr2 (Absmeier et al., 2015a). To gain insight into the ATPase mechanism of Brr2, we determined crystal structures of the cBrr2^T4^ variant in the nucleotide-free (apo) state, as well as in complex with ADP and the non-hydrolyzeable ATP analog, ATPγS, at resolutions ranging from 2.8 to 3.3 Å (Table 1). To achieve full nucleotide occupancy and to unequivocally locate coordinated divalent metal ions, cBrr2^T4^ was co-crystallized with ADP•AlF_3_ or ATPγS in the presence of Mn^2+^ ions, and the nucleotides and Mn^2+^ were included in cryo-protection buffers during crystal harvest.

**Table 1.** Crystallographic Data.

| Data collection |  |  |  |
| --- | --- | --- | --- |
| Structure | Apo | ADP | ATPyS |
| Wavelength [Å] | 0.9184 | 0.9184 | 1.8814 |
| Space group | P3<sub>1</sub> | P3<sub>1</sub> | P3<sub>1</sub> |
| Unit cell parameters |  |  |  |
| a (=b) [Å] | 124.5 | 125.1 | 124.8 |
| c [Å] | 128.0 | 127.4 | 127.1 |
| Resolution [Å]<sup>a</sup> | 50.0-3.3 (3.39-3.30) | 50.0-2.8 (2.97-2.80) | 50.0-2.7 (2.77-2.70) |
| Reflections |  |  |  |
| Total | 146,127 (10,060) | 579,441 (92,330) | 625,210 (45,316) |
| Unique | 33,329 (2,454) | 54,996 (8,834) | 60,260 (4,413) |
| Multiplicity | 4.4 (4.1) | 10.5 (10.5) | 10.4 (10.3) |
| Completeness [%] | 99.8 (99.7) | 99.5 (98.8) | 99.1 (98.1) |
| Mean I/σ(I) | 7.4 (1.2) | 13.6 (1.0) | 13.4 (1.0) |
| R<sub>meas</sub>(I) [%]<sup>b</sup> | 27.3 (157.6) | 21.0 (264.3) | 15.1 (251.6) |
| CC<sub>1/2</sub> [%]<sup>c</sup> | 98.1 (36.8) | 99.7 (43.1) | 99.9 (38.7) |
| Refinement |  |  |  |
| Structure | Apo | ADP | ATPyS |
| Resolution [Å]<sup>a</sup> | 50-3.3 (3.40-3.30) | 50-2.9 (3.00-2.90) | 50-2.8 (2.90-2.80) |
| Reflections |  |  |  |
| Unique | 33,295 (3,328) | 49,300 (4,891) | 54,133 (5,401) |
| Test set [%] | 5.9 | 5 | 5 |
| R<sub>work</sub> [%]<sup>d</sup> | 20.6 (32.2) | 22.0 (35.3) | 20.8 (30.9) |
| R<sub>free</sub> [%]<sup>e</sup> | 26.2 (37.6) | 27.1 (42.1) | 25.9 (36.3) |
| Contents of A.U.<sup>(f)</sup> |  |  |  |
| Non-H atoms | 13,495 | 13,417 | 13,506 |
| Protein residues | 1680 | 1657 | 1666 |
| ADP/ATPyS | 0 | 3 | 3 |
| Mn<sup>2+</sup> ions | 0 | 2 | 3 |
| Water oxygens | 2 | 43 | 40 |
| Mean B factors [Å<sup>2</sup>] |  |  |  |
| Wilson | 77.4 | 71.5 | 73.4 |
| Model atoms | 89.5 | 89.2 | 91.2 |
| Rmsd<sup>(g)</sup> from ideal geometry |  |  |  |
| Bond lengths [Å] | 0.003 | 0.003 | 0.002 |
| Bond angles [°] | 0.58 | 0.61 | 0.53 |
| Model quality<sup>(h)</sup> |  |  |  |
| Overall score | 2.30 | 2.28 | 1.96 |
| Clash score | 9.95 | 8.40 | 6.42 |
| Ramachandran favored [%] | 95.9 | 96.3 | 97.1 |
| Ramachandran outliers [%] | 0.0 | 0.0 | 0.0 |
| PDB ID | 6QWS | 6QV3 | 6QV4 |

<sup>a</sup> Values in parentheses refer to the highest resolution shells.
<sup>b</sup> $R_{meas}(I) = \sum_h [N/(N-1)]^{1/2} \sum_i |I_{ih} - \langle I_h \rangle| / \sum_h \sum_i I_{ih}$ , in which $\langle I_h \rangle$ is the mean intensity of symmetry-equivalent reflections $h$ , $I_{ih}$ is the intensity of a particular observation of $h$ and $N$ is the number of redundant observations of reflection $h$ .
<sup>c</sup> $CC_{1/2} = (\langle I^2 \rangle - \langle |I|^2 \rangle) / (\langle I^2 \rangle - \langle |I|^2 \rangle + \sigma_\epsilon^2)$ , in which $\sigma_\epsilon^2$ is the mean error within a half-dataset (Karplus and Diederichs, 2015).
<sup>d</sup> $R_{work} = \sum_h |F_o - F_c| / \sum F_o$ (working set, no $\sigma$ cut-off applied).
<sup>e</sup> $R_{free}$ is the same as $R_{work}$ , but calculated on the test set of reflections excluded from refinement.
<sup>f</sup> A.U. – asymmetric unit. <sup>g</sup> Rmsd – root-mean-square deviation <sup>h</sup> Calculated with MolProbity (Chen et al., 2010).

The overall structure of cBrr2^T4^ without or with nucleotides resembles previously reported Brr2 helicase structures (Mozaffari-Jovin et al., 2013; Nguyen et al., 2013; Santos et al., 2012) (Absmeier 2016) (Figure 1B). We did not observe density for the very N-terminal residues (apo cBrr2^T4^, residues 473-478; cBrr2^T4^-ADP, residues 473-481; cBrr2^T4^-ATPγS, residues 473-477). Additionally, one loop in the RecA2 domain of the NC (cBrr2^T4^-ADP, residues 741-747; cBrr2^T4^-ATPγS, residues 743-746) and parts of the C-terminal IG domain (apo cBrr2^T4^, residues 2083-2093, 2105-2111, 2138-2146, 2158-2163, 2186-2193; cBrr2^T4^-ADP, residues 2080-2093, 2105-2111, 2160-2170, 2184-2193; cBrr2^T4^-ATPγS, residues 2077-2093, 2105-2110, 2149-2150, 2160-2170, 2184-2193) could not be built due to missing electron density. Upon co-crystallization with ADP•AlF_3_ or ATPγS, we observed ADP and ATPγS, respectively, in the canonical binding pockets between the RecA1 and RecA2 domains of the NC and the CC (Figure 1B,C). No density for the AlF_3_ moiety was observed upon co-crystallization with ADP•AlF_3_. Unexpectedly, a third ADP or ATPγS nucleotide bound between the two cassettes (Figure 1B,C). In the apo cBrr2^T4^ structure, sulfate ions are bound at motif I of the RecA1 domains of the NC and CC, and an additional sulfate ion is bound between the cassettes (Figure 1C). The sulfate ion in the ATPase-active nucleotide binding pocket of the NC may resemble a phosphate after nucleotide hydrolysis, as observed for other helicases (Schmitt et al., 2018). Electron densities of the respective nucleotides and the sulfate ions were well defined in both helicase cassettes and in between the cassettes in the apo cBrr2^T4^, cBrr2^T4^-ADP and cBrr2^T4^-ATPγS structures (Figure 1C). Positions of divalent cations were unequivocally derived from the anomalous signals of Mn^2+^ ions (Figure 1C).

### Nucleotide Binding to the Active NC Induces Conformational Changes

By truncating residues 1-425, we have previously produced the cBrr2^T3^ variant, which retains a portion of the NTR (the NC-clamp) that encircles the NC and partially inhibits the ATPase and helicase activities (Absmeier et al., 2015a). Moreover, we have previously determined a crystal structure of cBrr2^T3^ in complex with the Jab1 domain of the cPrp8 protein; cBrr2^T3^ could not be crystallized in the absence of cJab1 (Absmeier et al., 2016a). Comparison of the current cBrr2^T4^ structures with the cBrr2^T3^-cJab1 complex structure revealed that the two RecA domains of the NC approach each other more closely in the apo cBrr2^T4^ structure compared to the cBrr2^T3^-cJab1 complex (distance of the Cα-atoms of G554 of motif I and G904 of motif VI 6 Å *vs*. 8 Å, respectively; Figure 2A,B). The relative positions of the NC RecA domains in the ADP-bound cBrr2^T4^ structure remain similar to those of the apo cBrr2^T4^ structure (Figure 2B,C). However, the RecA1 and RecA2 domains of the NC further approach each other upon ATPγS binding (Cα-atom of G554 of motif I and Cα-atom of G904 of motif VI spaced 2 Å closer than in the apo and ADP-bound structures, and 4 Å closer than in the cBrr2^T3^-cJab1 complex structure; Figure 2D). The observed closing-in of motifs I and VI on the nucleotide is consistent with the idea that these elements need to contact bound ATP to facilitate ATP hydrolysis (Fairman-Williams et al., 2010). Consistently, similar situations have been observed in other RecA domain-containing ATPases (Pyle, 2008).

**Figure 2.**
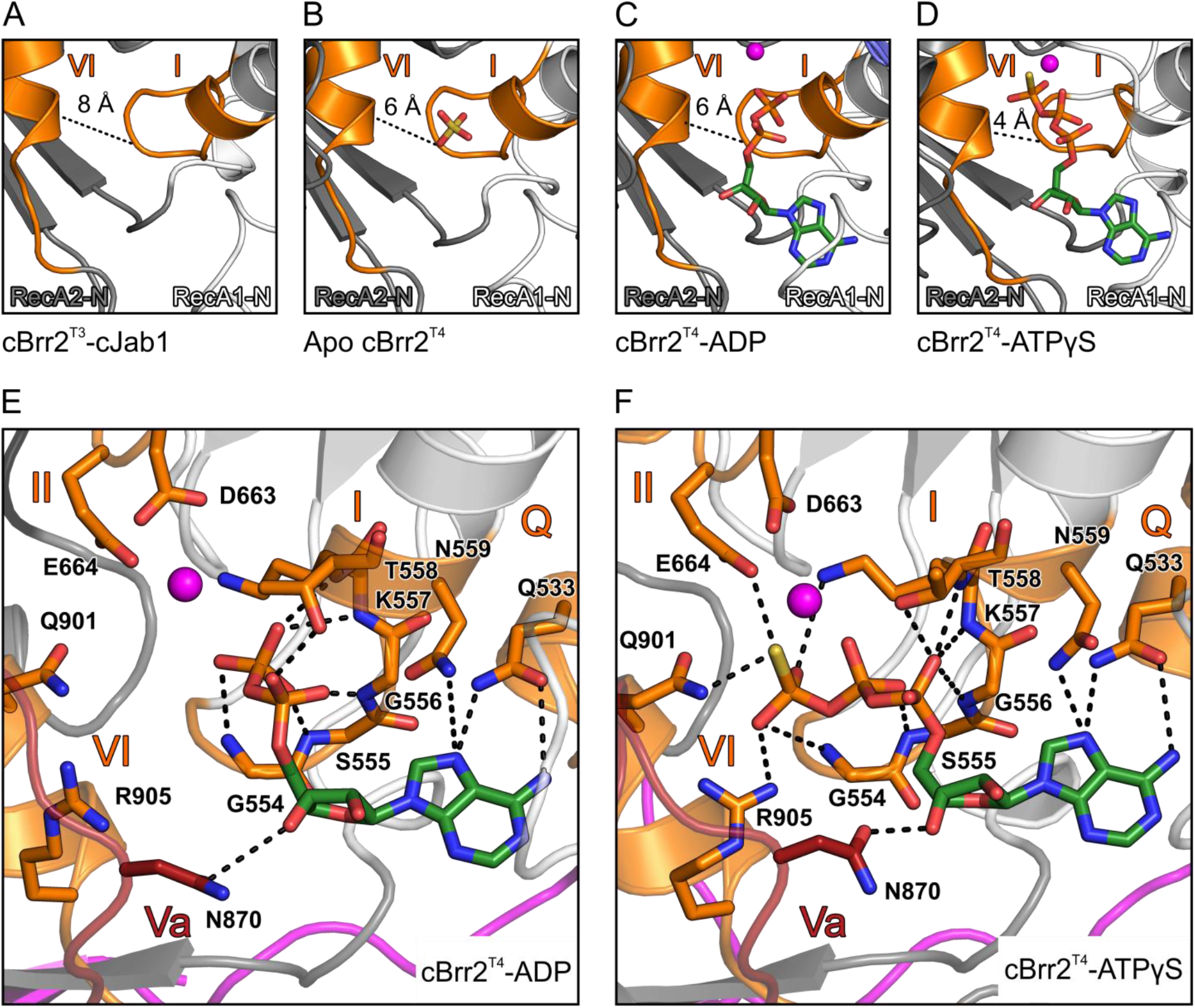
Nucleotide-Induced Conformational Changes in the NC. (A-D) Comparison of the distances (dashed lines) between motif I (Cα-atom of G554 in RecA1-N) and motif VI (Cα-atom of G904 in RecA2-N) in cBrr2^T3^-cJab1 (A; cJab1 has been omitted for clarity), apo cBrr2^T4^ (B), cBrr2^T4^-ADP (C) and cBrr2^T4^-ATPγS (D). RecA1/2-N, RecA1/2 domains of NC. Domain coloring as in Figure 1; motif I and motif VI, orange. Sulfate ions, ADP and ATPγS are shown as sticks and colored by atom type (carbon, dark green; nitrogen, blue; oxygen, red; phosphorus, orange; sulfur, yellow). Magenta spheres, Mn^2+^ ions. (E,F) Details of the nucleotide coordination in the NC of cBrr2^T4^-ADP (E) and cBrr2^T4^-ATPγS (F). Coloring of domains/regions, nucleotides and Mn^2+^ ions as in (A-D). Interacting residues are shown as sticks and colored by atom type (carbon as the respective domain/region; nitrogen, blue; oxygen, red). For interactions involving only protein backbone atoms, side chains are not shown for clarity. Dashed lines indicate hydrogen bonds or salt bridges. Structures were aligned with respect to their motifs I in the NC RecA1 domains.

In both nucleotide-bound cBrr2^T4^ structures, the conserved Q-motif of the NC (Q533) coordinates the adenine base through hydrogen bonding at positions N6 and N7 (Figure 2E,F). Conserved residues of motif I (555-559) interact with the phosphates of ADP and ATPγS (Figure 2E,F). N559 additionally contacts the adenine base. N870 (motif Va; important for coupling of ATP hydrolysis to RNA duplex unwinding) binds the ribose moieties of the nucleotides. The γ-thiophosphate of ATPγS is additionally recognized by E664 (motif II), Q901 (motif VI) and R905 (motif VI) in the cBrr2^T4^-ATPγS structure, while in the ADP-bound structure residues of motif II do not contact the nucleotide (Figure 2E,F).

The above comparisons suggest that the NC-clamp (contained in cBrr2^T3^ but absent from cBrr2^T4^) stabilizes the RecA domains of the NC in an open, NTPase-inactive conformation, explaining the previously observed lower intrinsic ATPase activity of cBrr2 truncations that include the NC-clamp compared to cBrr2^T4^ (Absmeier et al., 2015a). They also illustrate how only ATP, but not ADP, can engage motifs II and VI, leading to further NTPase-supporting closure of the nucleotide binding cleft.

### The Intrinsic ATPase Activity of cBrr2^T4^ Is Attenuated by Exclusion of an Attacking Water

Comparison of our cBrr2^T4^-ATPγS structure with the *C. thermophilum* DEAH/RHA-box RNA helicase Prp43 (cPrp43) bound to ADP•BeF_3_, mimicking bound ATP in the ground state (Tauchert et al., 2017), revealed a strikingly different position of a conserved glutamine (Q901 in cBrr2; Q428 in cPrp43) and arginine (R908 in cBrr2; R435 in cPrp43) in motif VI relative to the respective ATP analog (Figure 3A,B). In the cPrp43 structure (Figure 3B), Q428 resides approximately 6 Å from the BeF_3_ (representing the γ-phosphate) and positions (and possibly polarizes), together with E219 (motif II) and R432 (motif VI), an attacking water at 3 Å distance from and in line with the “scissile” bond of the γ-phosphate surrogate. Additionally, R435 of motif VI (arginine finger) interacts with the β-phosphate and BeF_3_ in the cPrp43 structure. In the cBrr2^T4^-ATPγS structure, the corresponding Q901 is positioned much closer (3 Å) to and engages in direct interactions with the γ-thiophosphate (Figure 3A). This close approach apparently prevents positioning of an attacking water molecule in the cBrr2^T4^-ATPγS structure (Figure 3A). In addition, R908 (arginine finger) is flipped away from the bound nucleotide and does not contact the β and γ-phosphates.

**Figure 3.**
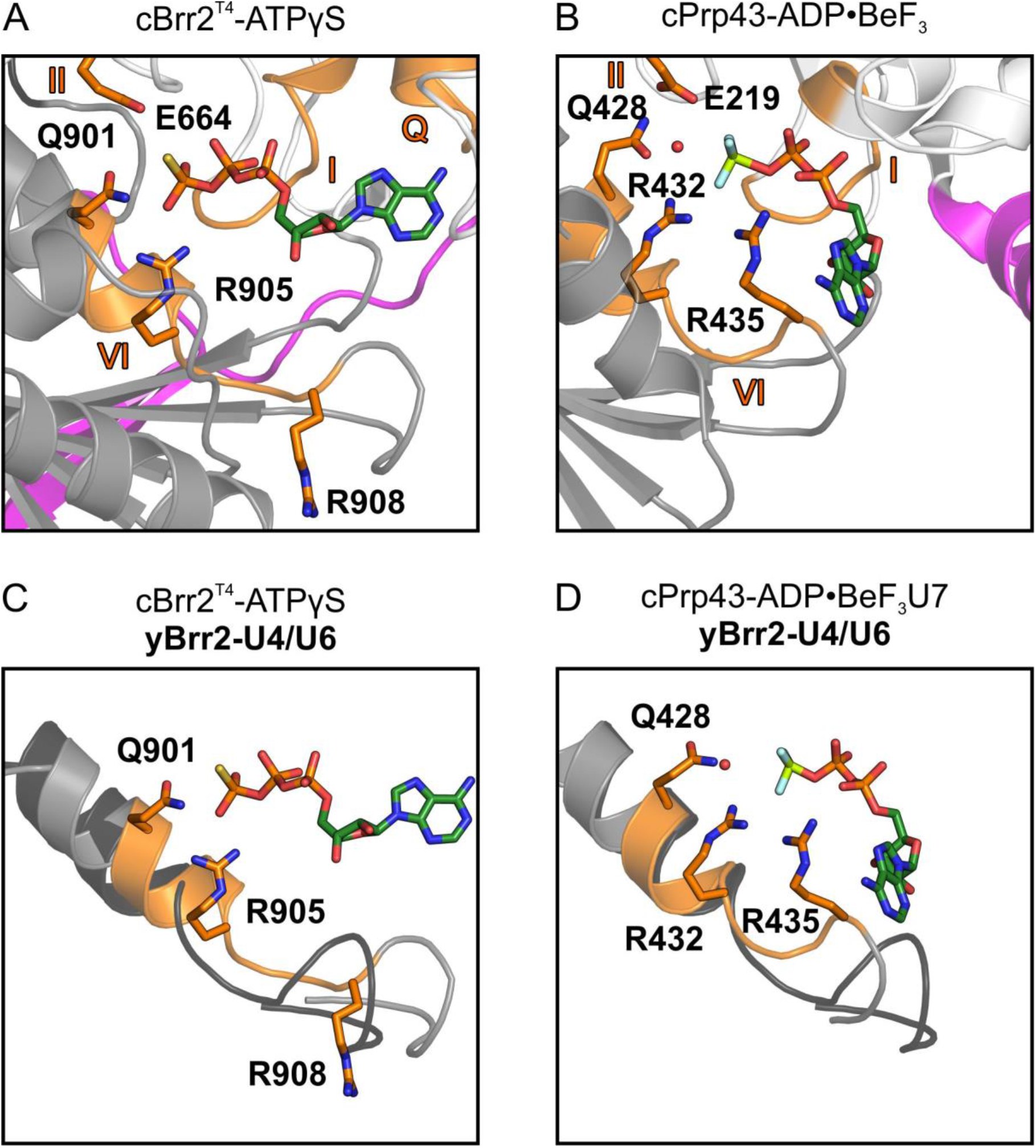
Comparison of ATP Analog Binding by cBrr2^T4^ and cPrp43. (A) ATPγS bound by Q901 and R905 in the NC of cBrr2^T4^ and position of E664 and R908 relative to the nucleotide. (B) ADP•BeF_3_ binding and positioning of an attacking water by E219, Q428 and R432 and position of R435 relative to the nucleotide in a cPrp43-ADP•BeF_3_ structure (PDB ID 5LTJ). (C) Superposition of cBrr2^T4^-ATPγS (cBrr2^T4^, gray and orange) and yeast (y) Brr2 (black) bound to U4/U6 di-snRNA from a yeast U4/U6•U5 tri-snRNP structure (PDB ID 5GAN). (D) Superposition of cPrp43-ADP•BeF_3_ (cPrp43, gray and orange) bound to a U7 RNA oligo (PDB ID 5LTA) and yBrr2 (black) bound to U4/U6 di-snRNA from a yeast U4/U6•U5 tri-snRNP structure (PDB ID 5GAN). Only the α-helices containing motifs VI are shown for clarity. Domain coloring as in Figure 1. Motifs Q, I, II and VI, orange. Residues from motifs II and VI are shown as sticks and colored by atom type (carbon, orange; nitrogen, blue; oxygen, red). Nucleotides are shown as sticks and colored by atom type (beryllium, light green; carbon, dark green; nitrogen, blue; oxygen, red; fluorine, light blue; phosphorus, orange; sulfur, yellow). Red sphere, water oxygen. Structures are aligned with respect to their motifs I in the NC RecA1 domains.

Superposition of the cBrr2^T4^-ATPγS structure with Brr2 bound to U4/U6 di-snRNA in a yeast U4/U6•U5 tri-snRNP (Nguyen et al., 2016) or pre-catalytic B complex (Bertram et al., 2017; Plaschka et al., 2018; Zhan et al., 2018) suggests a movement of the helix bearing motif VI, and thus Q901 and R908, upon RNA binding (Figure 3C). This conformational change may displace Q901 from the γ-phosphate and allow it to productively position an attacking water, as seen in the cPrp43-ADP•BeF_3_ structure. In addition, the movement of motif VI would bring R908 closer to the ATP β and γ-phosphates, where it can contribute to nucleotide hydrolysis (Figure 3C). Superposition of cPrp43 bound to RNA and ADP•BeF_3_ and yeast Brr2 bound to U4/U6 di-snRNA reveals a similar position of motif VI, further supporting this model (Figure 3D). Thus, in addition to NC-clamp-mediated RecA domain opening, exclusion of an attacking water molecule by direct interaction of Q901 and the ATP γ-phosphate in the absence of substrate RNA appears to be another mechanism that ensures a low intrinsic ATPase activity in Brr2.

### Nucleotide Binding at the CC and Cassette Interface

Unlike in the active NC, the RecA1 and RecA2 domains of the CC do not approach each other upon ATPγS binding (Figure S1A,B). While a similar situation has previously also been observed for an N-terminal truncation variant of human Brr2 (Santos et al., 2012), in this latter case nucleotides could only be soaked into pre-formed apo crystals after chemical crosslinking, leaving the authentic conformational response of the CC to nucleotide binding an open question.

Interestingly, in both nucleotide-bound cBrr2^T4^ structures, an additional ADP or ATPγS molecule was observed, wedged between the RecA1 and IG domains of the NC and the RecA2 and HB domains of the CC (Figure S1C). This ADP/ATPγS is mainly coordinated by side chain interactions with the phosphates. In addition, there are two main chain interactions with the base and one with the ribose (Figure S1C). The additional nucleotide binding site does not resemble any of the typical nucleotide binding pockets known in RNA helicases. Superposition of cBrr2^T4^-ATPγS and yeast Brr2 bound to U4/U6 di-snRNA revealed that the additional nucleotide binding site represents part of the U4/U6 di-snRNA binding surface of Brr2 (Figure S1D). A nucleotide bound at this position may thus interfere with RNA binding and/or may restrict cassette movements possibly required for RNA duplex unwinding. To test these ideas, we performed gel-based unwinding assays with increasing concentrations of ATP. In agreement with the structures, we observed inhibition of cBrr2^T4^-mediated U4/U6 unwinding at very high ATP concentrations (15 mM; Figure S1E). Recently, a Brr2-specific small-molecule inhibitor was identified, which binds at the interface of the two helicase cassette, but at a site different from the additional nucleotide binding site in the present structures (Iwatani-Yoshihara et al., 2017). Thus, our structural results reveal a new binding pocket at the cassette interface of Brr2, which may be further exploited for inhibitor design.

## DISCUSSION

### Influence of N-Terminal Regions on Nucleotide Binding in Ski2-Like Helicases

Here, we analyzed the structural basis of nucleotide binding to the Ski2-like RNA helicase, Brr2. Our results revealed multiple levels of auto-inhibition that prevent futile intrinsic ATPase activity. To date only two other structures of Ski2-like helicases bound to an ATP analog have been reported. In the AMPPNP-bound structure of yeast Ski2 (Halbach et al., 2012), the two RecA domains are in close proximity and the AMPPNP phosphates are recognized by motifs I and VI, resembling the cBrr2^T4^-ATPγS structure. However, due to the lack of divalent cations in the crystallization condition, the density for the γ-phosphate is weakly defined in the Ski2-AMPPNP structure, and not all hydrolysis-relevant contacts to the nucleotide are formed. As in the present nucleotide-bound structure of cBrr2^T4^, the Ski2 arginine finger of motif VI is flipped away and does not contact the nucleotide, suggesting that Ski2-like RNA helicases in general bind ATP initially in a pre-hydrolysis state. Interestingly, Ski2 also contains a long NTR that was removed for crystallization. As in the case of Brr2, removal of the NTR might have led to a movement of the RecA domains towards each other; thus, modulation of nucleotide binding/hydrolysis by an N-terminal region may be a recurring theme in some Ski2-like enzymes.

In contrast, no conformational changes were observed in the *Pyrococcus furiosus* Hel308 protein, Hjm, upon AMPPCP binding (Oyama et al., 2009). The ATP analog interacted only with motifs located in the RecA1 domain (Oyama et al., 2009). Moreover, in the Hjm-AMPPCP structure neither of the conserved arginines in motif VI engages in interactions with the nucleotide, supporting the idea that an inactive state has been captured. Hjm contains a very short NTR compared to Brr2 and Ski2. This NTR folds back onto the RecA1 domain but does not concomitantly contact the RecA2 domain and, thus, is not expected to restrict RecA domain conformational changes required for stable nucleotide binding. A possible explanation for the lack of conformational changes upon AMPPCP binding in this study may be that the nucleotide bound-state of Hjm was obtained upon soaking of apo Hjm crystals with AMPPCP. The two RecA domains engage in several crystal contacts, which might have prevented them from adopting a closed conformation upon AMPPCP binding. In any case, the analysis illustrates that NTR-based modulation of nucleotide binding is apparently not universally conserved in Ski2-like helicases.

### RNA Binding Triggers Adoption of an ATPase-Competent State

Our structures of apo cBrr2^T4^, cBrr2^T4^-ADP and cBrr2^T4^-ATPγS reveal a movement of the RecA1 and RecA2 domains towards each other upon ATPγS, but not ADP, binding, as observed for other ATPases containing two RecA domains (Ye et al., 2004). ATPγS is highly coordinated in our cBrr2^T4^-ATPγS structure by motifs of both RecA domains, resembling ADP-BeF_3_ in the active site of the cPrp43 DEAH/RHA-box helicase (Tauchert et al., 2017). However, there are important differences. The arginine finger residing in motif VI (R908) does not contact the nucleotide in the cBrr2^T4^-ATPγS structure, which would be necessary for a fully hydrolysis-competent conformation. In addition, there is no evidence for a water molecule positioned in line with the β-γ phospho-anhydride bond of ATPγS (representing the scissile bond of ATP). Comparison to the cPrp43-ADP•BeF_3_ structure (Tauchert et al., 2017) indicates that a conserved glutamine of motif VI in Brr2 has to be relocated to allow positioning of an attacking water (Figure 3A). We suggest that RNA binding is the final trigger that induces a switch in Brr2 towards the fully hydrolysis-competent conformation. Such a mechanism would additionally aid in preventing unproductive ATP hydrolysis in the absence of RNA. In full agreement with this idea, the conformation of Brr2, when bound to its U4/U6 substrate (Bertram et al., 2017; Nguyen et al., 2016; Plaschka et al., 2018; Zhan et al., 2018) reveals conformational rearrangements towards a hydrolysis-competent state (Figure 3C).

### Helicases Exhibit Family-Specific Nucleotide Hydrolysis Mechanisms

The structures of cPrp43 bound to ADP and ADP•BeF_3_ (Tauchert et al., 2017) showed surprising flexibility of certain domains upon binding to different nucleotides (Figure 4A). Binding of ADP•BeF_3_ results in a rotation of the RecA2 domain, which breaks the interactions of a long β hairpin in the RecA2 domain (the equivalent of the separator loop [SL] in Ski2-like helicases), with the HB domain (Figure 4A). As a consequence, the HB/ratchet and OB-fold domains, which form part of the RNA binding tunnel, open up and form a groove, which can accommodate even complex RNA folds. Superposition of the apo cBrr2^T4^ and cBrr2^T4^-ATPγS structures, reveal that the SL of Brr2 still interacts with the HB domain and that the HB/HLH/IG domains (the Sec63 homology region) rotate together with the RecA2 domain upon ATPγS binding (Figure 4B), similar to the situation in Ski2 (Halbach et al., 2012). In agreement with the different behavior upon nucleotide binding, the SL-like elements has different functions in different SF2 members. The elongated SL-like element in cPrp43 is thought to control access to the RNA binding site (Tauchert et al., 2017). In Ski2-like helicases, the SL is much shorter and acts as a tool that separates the strands of the substrate RNA duplex (Büttner et al., 2007; Woodman et al., 2007). In addition, unlike in our cBrr2^T4^ structures, an attacking water molecule is already present in cPrp43 in the absence of RNA, and RNA binding does not significantly change the ADP•BeF_3_ coordination.

**Figure 4.**
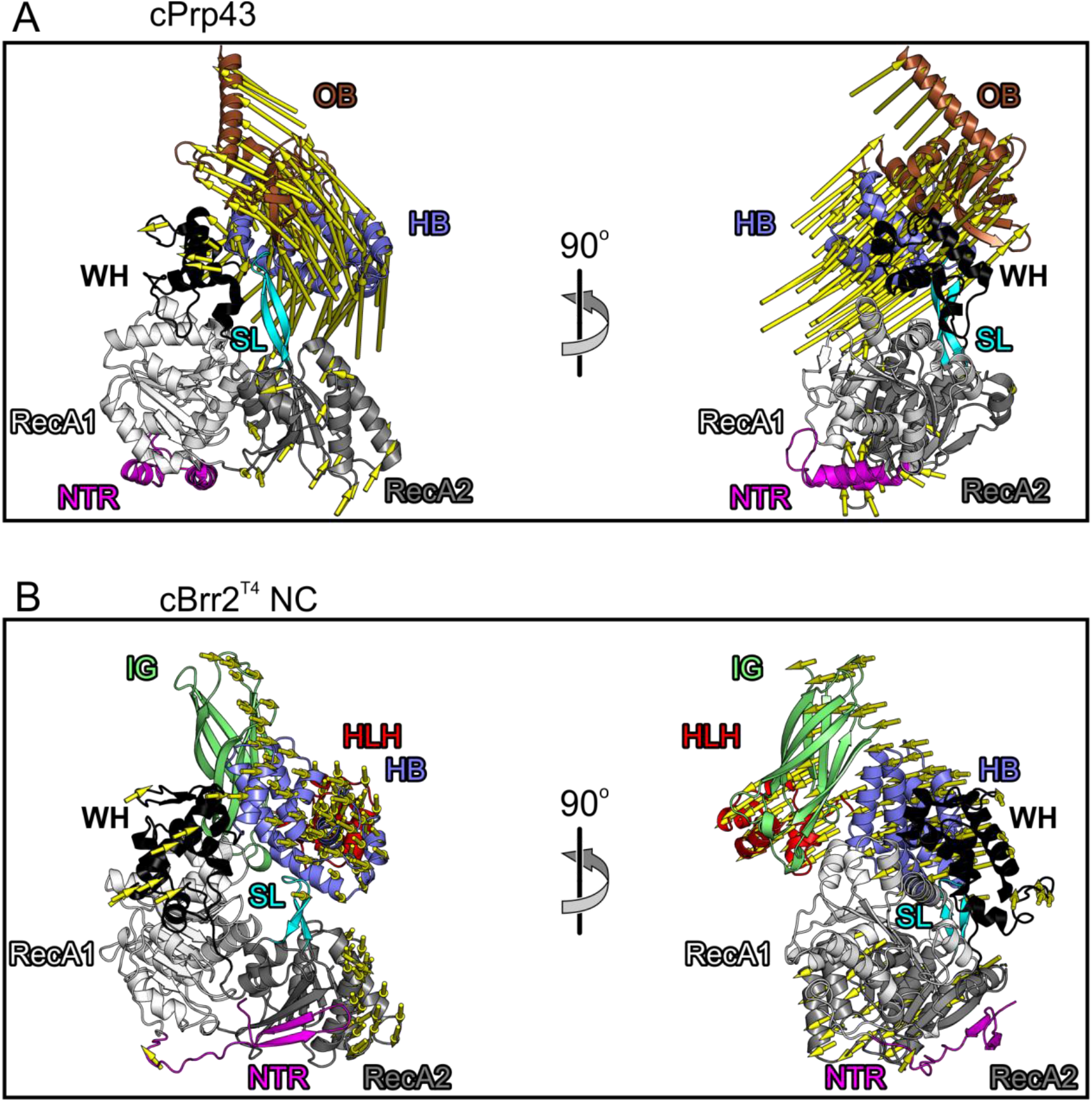
Domain Movements in cPrp43 and cBrr2^T4^ Upon ATP Analog Binding. (A) Domain movements in cPpr43 from the ADP-bound conformation (base of mode vectors; PDB ID 5D0U) to the ADP•BeF_3_-bound conformation (head of mode vectors; PDB ID 5LTJ). (B) Domain movements in the cBrr2^T4^ NC from the ADP-bound conformation (base of mode vectors) to the ATPγS-bound conformation (head of mode vectors). OB, brown; separator loop (SL), cyan; other domains/regions are colored as in Figure 1. Structures were aligned with respect to their motifs I in the (NC) RecA1 domains. cPrp43 and cBrr2^T4^ NC are shown with the RecA domains in the same orientation.

Taken together, the mechanism of RecA domain closure and coordination of the attacking water by a glutamate in motif II and by glutamine and arginine residues in motif VI may be conserved, but the trigger for adopting the ultimate hydrolysis-competent state seems to differ in different SF2 families. In the future, further structural studies of Ski-2 like helicases bound to nucleic acid and nucleotides need to be performed to elucidate the exact mechanism of ATP hydrolysis and to understand the differences to other SF2 families.

## SUPPLEMENTAL INFORMATION

Supplemental Information includes Figure and can be found with this article online at https://…

## ACKNOWLEDGEMENTS

We acknowledge access to beamline BL14.1 of the BESSY II storage ring (Berlin, Germany) *via* the Joint Berlin MX Laboratory sponsored by Helmholtz Zentrum Berlin für Materialien und Energie, Freie Universität Berlin, Humboldt-Universität zu Berlin, Max-Delbrück Centrum, Leibniz-Institut für Molekulare Pharmakologie and Charité – Universitätsmedizin Berlin. We thank Dr. Manfred S. Weiss for support during diffraction data collection. This work was supported by a grant from the Deutsche Forschungsgemeinschaft (TRR186-A15) to MCW. KFS was supported by a Dahlem International PostDoc Fellowship of Freie Universität Berlin.

## AUTHOR CONTRIBUTIONS

EA purified protein and performed activity assays and crystallization trials, determined and refined crystal structures and wrote the manuscript with KFS and MCW. All authors participated in data interpretation.

## STAR METHODS

### CONTACT FOR REAGENT AND RESOURCE SHARING

Requests for resources and reagents should be directed to and will be fulfilled by the Lead Contact, Markus C. Wahl.

### METHOD DETAILS

#### Protein Production and Purification

A codon-optimized DNA fragment encoding residues 473-2193 of *C. thermophilum* Brr2 was cloned into a modified pFL vector (EMBL, Grenoble) and verified by sequencing to produce cBrr2^T4^ bearing a TEV protease-cleavable N-terminal His10-tag (Absmeier et al., 2015a). The plasmid was transformed into *E. coli* DH10MultiBacY cells and further integrated *via* Tn7 transposition into the baculovirus genome (EMBacY) maintained as a bacterial artificial chromosome (BAC) (Trowitzsch et al., 2010). The Tn7 transposition site was embedded in a *lacZα* gene allowing the selection of positive EMBacY recombinants *via* blue/white screening. Recombinant EMBacY was isolated from the bacterial host and used to transfect Sf9 cells (Invitrogen).

For initial virus (V_0_) production, recombinant EMBacY was transfected into adhesive Sf9 cells (Invitrogen) in 6-well plates. The efficiency of transfection was monitored by eYFP fluorescence. The V0 virus generation was used to infect 50 ml Sf9 cells for virus amplification. The second, high titer virus generation (V1) was then used to infect 1200 ml High Five™ cells (Invitrogen) for large scale protein production (Santos et al., 2012). The infected cells were harvested when the eYFP signal reached a plateau and before the cell viability dropped below 90 %.

The High Five^™^ cell pellet was resuspended in 40 mM HEPES-NaOH, pH 8.0, 600 mM NaCl, 1 mM DTT, 1.5 mM MgCl2, 20 mM imidazole, supplemented with EDTA-free protease inhibitor (Roche) and lyzed by sonication using a Sonopuls Ultrasonic Homogenizer HD 3100 (Bandelin). The target protein was captured from the cleared lysate on a 5 ml HisTrap FF column (GE Healthcare) and eluted with a linear gradient from 20 to 500 mM imidazole. The His-tag was cleaved with TEV protease during overnight dialysis at 4 °C against 40 mM HEPES-NaOH, pH 8.0, 600 mM NaCl, 1 mM DTT. The cleaved protein was again loaded on a 5 ml HisTrap FF column to remove the His-tagged protease, uncut protein and cleaved His-tag. The flow-through containing the protein of interest was diluted to a final concentration of 80 mM NaCl, treated with RNaseA and loaded on a 5 ml Heparin column (GE Healthcare) equilibrated with 40 mM HEPES-NaOH, pH 8.0, 50 mM NaCl, 1 mM DTT. The protein was eluted with a linear 0.05 to 1.5 M NaCl gradient and further purified by gel filtration on a 16/60 Superdex 200 gel filtration column (GE Healthcare) in 10 mM Tris-HCl, pH 8.0, 200 mM NaCl, 2 mM DTT. Protein for activity assays was purified similarly, except that 10, 5 and 20 (v/v) % glycerol was added to the buffer for the HisTrap, Heparin and gel filtration step, respectively, and that the His-tag was not cleaved.

#### Crystallographic Procedures

Fractions containing the target protein were pooled, concentrated to 12 mg/ml and used for crystallization. Apo cBrr2^T4^ crystals were grown in 24-well plates using the sitting-drop vapor diffusion technique at 18 °C with drops containing 1 μl protein complex solution and 1 μl reservoir solution (0.1 M Tris-HCl, pH 8.0, 24 % (w/v) PEG 3350, 0.2 M LiSO_4_). Initial crystals were further optimized by seeding. Crystals were cryo-protected by transfer into mother liquor containing 25 % (v/v) ethylene glycol and flash-cooled in liquid nitrogen. ADP- and ATPγS-bound cBrr2^T4^ crystals were grown in 24-well plates using the sitting-drop vapor diffusion technique at 18 °C with drops containing 1 μl protein solution (9 mg/ml; supplemented with 2 mM ADP•AlF_3_/ATPγS and 6 mM MgCl_2_) and 1 μl reservoir solution (0.1 M Tris-HCl, pH 8.0, 9 % (w/v) PEG 8000, 0.2 M Ca(OAc)_2_) and further optimized by seeding. Crystals were soaked and cryo-protected by transfer into mother liquor supplemented with the respective nucleotides and cryo-protectant (0.1 M Tris-HCl, pH 8.0, 9 % (w/v) PEG 8000, 25 % (v/v) PEG 400, 0.2 M NaOAc, 12 mM ADP•AlF_3_/6 mM MnCl_2_ or 25 mM ATPγS/12 mM MnCl_2_) and flash-cooled in liquid nitrogen.

Diffraction data were collected at 100 K on beamline 14.1 of the BESSY II storage ring, Berlin, Germany (Mueller et al., 2015) with a monochromated X-ray beam (λ = 0.9184 Å or 1.8814 Å) and processed with XDSAPP (Sparta et al., 2016) (Table 1). The structures were solved by molecular replacement with PHASER (McCoy et al., 2007) using cBrr2^T3^-cJab1 structure coordinates as a search model (PDB ID 5M59) (Absmeier et al., 2016a). The structures were refined by alternating rounds of manual model building with Coot (Emsley et al., 2010) and automated refinement with PHENIX (Adams et al., 2002; Afonine et al., 2012) (Table 1).

#### Structural Comparisons

Structures were superimposed using the Secondary Structure Matching tool of Coot.

#### Unwinding Assays

Unwinding assays were conducted and evaluated as described (Mozaffari-Jovin et al., 2013; Santos et al., 2012). Briefly, U4/U6 di-snRNA complex (2 nM) and cBrr2^T4^ (100 nM) were mixed in 40 mM Tris-HCl, pH 7.5, 50 mM NaCl, 1.5 mM DTT, 0.1 mg/ml acetylated BSA. After incubation for 3 min at 30 °C, reactions were started by the addition of different concentrations of ATP/MgCl2 (2, 4, 6, 8, 10, 25 mM). 10 μl samples were taken after 20 min, mixed with 10 μl 40 mM Tris-HCl, pH 7.4, 50 mM NaCl, 25 mM EDTA, 1 % (w/v) SDS, 10 % (v/v) glycerol, 0.05 % (w/v) xylene cyanol, 0.05 % (w/v) bromophenol blue and separated using 6 % RNA native PAGE (19:1). Gels were scanned on a phosphoimager.

### QUANTIFICATION AND STATISTICAL ANALYSIS

Not applicable.

### DATA AND SOFTWARE AVAILABILITY

Coordinates and structure factors for the cBrr2^T4^ apo and nucleotide-bound structures have been deposited in the Protein Data Bank (www.pdb.org) under access codes 6QWS (apo cBrr2^T4^), 6QV3 (cBrr2^T4^-ADP) and 6QV4 (cBrr2^T4^-ATPγS) and will be released upon publication.

## SUPPLEMENTAL FIGURES

**Figure S1.**
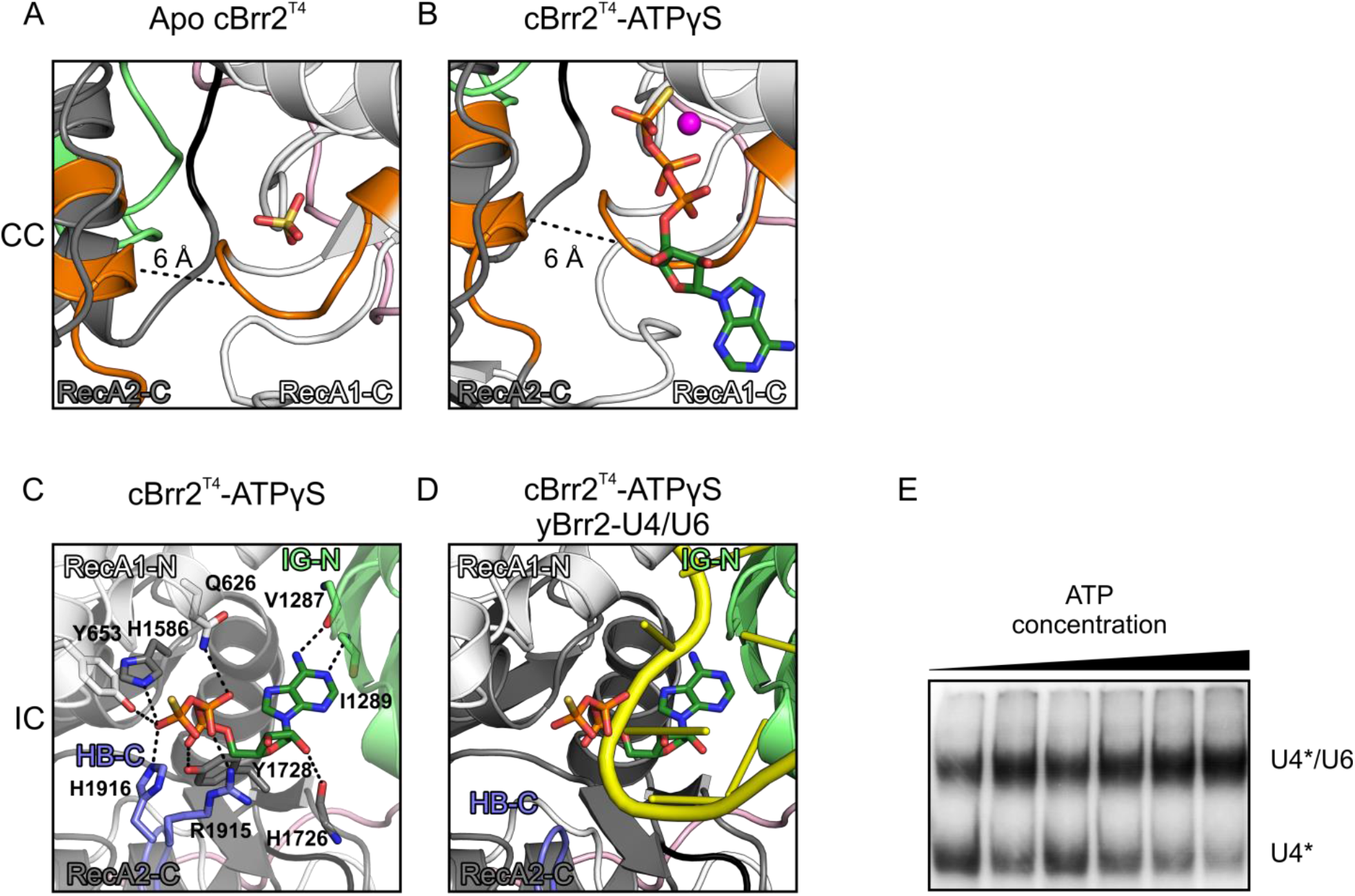
Nucleotide binding to the CC and between cassettes, related to Figure 2. (A,B) Distances between motif I (RecA1-C, Cα-atom of G1404) and motif VI (RecA2-C, Cα-atom of G1741) in the CCs of the apo cBrr2^T4^ (A) and cBrr2^T4^-ATPγS (B) structures. Coloring as in Figure 2. RecA1/2-C, RecA1/2 domains of the CC. (C) ATPγS binding between the cassettes by residues of the RecA1-N, IG-N, RecA2-C and HB-C domains. Domains are colored as in Figure 1 and interacting residues are shown as sticks and colored by atom type, as in Figure 2. IC, inter-cassette binding site; IG-N, immunoglobulin-like domain of the NC; HB-C, helical bundle domain of the CC. (D) Superposition of the cBrr2^T4^-ATPγS structure and yeast U4 snRNA from a yeast U4/U6•U5 tri-snRNP structure (PDB ID 5GAN). Domains and ATPγS are colored as in Figures 1 and 2 and RNA is colored yellow. (E) cBrr2^T4^-mediated U4/U6 di-snRNA unwinding (single time points) at increasing ATP concentrations (2, 4, 6, 8, 10, 25 mM ATP).

## REFERENCES

Absmeier, E., Becke, C., Wollenhaupt, J., Santos, K.F., and Wahl, M.C. (2016a). Interplay of cis- and trans-regulatory mechanisms in the spliceosomal RNA helicase Brr2. Cell Cycle.

Absmeier, E., Wollenhaupt, J., Mozaffari-Jovin, S., Becke, C., Lee, C.-T., Preussner, M., Heyd, F., Urlaub, H., Lührmann, R., Santos, K.F., et al. (2015a). The large N-terminal region of the Brr2 RNA helicase guides productive spliceosome activation. Genes Dev. 29, 2576–2587.

Absmeier, E., Rosenberger, L., Apelt, L., Becke, C., Santos, K.F., Stelzl, U., and Wahl, M.C. (2015b). A noncanonical PWI domain in the N-terminal helicase-associated region of the spliceosomal Brr2 protein. Acta Crystallogr. D Biol. Crystallogr. 71, 762–771.

Agafonov, D.E., Kastner, B., Dybkov, O., Hofele, R.V., Liu, W.-T., Urlaub, H., Lührmann, R., and Stark, H. (2016). Molecular architecture of the human U4/U6.U5 tri-snRNP. Science 351, 1416–1420.

Bertram, K., Agafonov, D.E., Dybkov, O., Haselbach, D., Leelaram, M.N., Will, C.L., Urlaub, H., Kastner, B., Lührmann, R., and Stark, H. (2017). Cryo-EM Structure of a Pre-catalytic Human Spliceosome Primed for Activation. Cell 170, 701–713.e11.

Büttner, K., Nehring, S., and Hopfner, K.-P. (2007). Structural basis for DNA duplex separation by a superfamily-2 helicase. Nat. Struct. Mol. Biol. 14, 647–652.

Chen, V.B., Arendall, W.B., Headd, J.J., Keedy, D.A., Immormino, R.M., Kapral, G.J., Murray, L.W., Richardson, J.S., and Richardson, D.C. (2010). MolProbity: all-atom structure validation for macromolecular crystallography. Acta Crystallogr. D Biol. Crystallogr. 66, 12–21.

Emsley, P., Lohkamp, B., Scott, W.G., and Cowtan, K. (2010). Features and development of Coot. Acta Crystallogr. D Biol. Crystallogr. 66, 486–501.

Fairman-Williams, M.E., Guenther, U.-P., and Jankowsky, E. (2010). SF1 and SF2 helicases: family matters. Curr. Opin. Struct. Biol. 20, 313–324.

Gu, M., and Rice, C.M. (2010). Three conformational snapshots of the hepatitis C virus NS3 helicase reveal a ratchet translocation mechanism. Proc. Natl. Acad. Sci. U. S. A. 107, 521–528.

Halbach, F., Rode, M., and Conti, E. (2012). The crystal structure of S. cerevisiae Ski2, a DExH helicase associated with the cytoplasmic functions of the exosome. RNA 18, 124–134.

Iwatani-Yoshihara, M., Ito, M., Klein, M.G., Yamamoto, T., Yonemori, K., Tanaka, T., Miwa, M., Morishita, D., Endo, S., Tjhen, R., et al. (2017). Discovery of Allosteric Inhibitors Targeting the Spliceosomal RNA Helicase Brr2. J. Med. Chem. 60, 5759–5771.

Jankowsky, E. (2011). RNA helicases at work: binding and rearranging. Trends Biochem. Sci. 36, 19–29.

Jankowsky, E., and Fairman, M.E. (2007). RNA helicases — one fold for many functions. Curr. Opin. Struct. Biol. 17, 316–324.

Karplus, P.A., and Diederichs, K. (2015). Assessing and maximizing data quality in macromolecular crystallography. Curr. Opin. Struct. Biol. 34, 60–68.

Kim, D.H., and Rossi, J.J. (1999). The first ATPase domain of the yeast 246-kDa protein is required for in vivo unwinding of the U4/U6 duplex. RNA 5, 959–971.

Laggerbauer, B., Achsel, T., and Lührmann, R. (1998). The human U5-200kD DEXH-box protein unwinds U4/U6 RNA duplices in vitro. Proc. Natl. Acad. Sci. 95, 4188–4192.

McCoy, A.J., Grosse-Kunstleve, R.W., Adams, P.D., Winn, M.D., Storoni, L.C., and Read, R.J. (2007). Phaser crystallographic software. J. Appl. Crystallogr. 40, 658–674.

Mozaffari-Jovin, S., Wandersleben, T., Santos, K.F., Will, C.L., Lührmann, R., and Wahl, M.C. (2013). Inhibition of RNA Helicase Brr2 by the C-Terminal Tail of the Spliceosomal Protein Prp8. Science 341, 80–84.

Mueller, U., Förster, R., Hellmig, M., Huschmann, F.U., Kastner, A., Malecki, P., Pühringer, S., Röwer, M., Sparta, K., Steffien, M., et al. (2015). The macromolecular crystallography beamlines at BESSY II of the Helmholtz-Zentrum Berlin: Current status and perspectives. Eur. Phys. J. Plus 130, 1–10.

Nguyen, T.H.D., Galej, W.P., Bai, X., Oubridge, C., Newman, A.J., Scheres, S.H.W., and Nagai, K. (2016). Cryo-EM structure of the yeast U4/U6.U5 tri-snRNP at 3.7 Å resolution. Nature 530, 298–302.

Oyama, T., Oka, H., Mayanagi, K., Shirai, T., Matoba, K., Fujikane, R., Ishino, Y., and Morikawa, K. (2009). Atomic structures and functional implications of the archaeal RecQ-like helicase Hjm. BMC Struct. Biol. 9, 2.

Plaschka, C., Lin, P.-C., Charenton, C., and Nagai, K. (2018). Prespliceosome structure provides insights into spliceosome assembly and regulation. Nature 559, 419–422.

Putnam, A.A., and Jankowsky, E. (2013). DEAD-box Helicases as Integrators of RNA, Nucleotide and Protein Binding. Biochim. Biophys. Acta 1829, 884–893.

Raghunathan, P.L., and Guthrie, C. (1998). RNA unwinding in U4/U6 snRNPs requires ATP hydrolysis and the DEIH-box splicing factor Brr2. Curr. Biol. 8, 847–855.

Santos, K.F., Jovin, S.M., Weber, G., Pena, V., Lührmann, R., and Wahl, M.C. (2012). Structural basis for functional cooperation between tandem helicase cassettes in Brr2-mediated remodeling of the spliceosome. Proc. Natl. Acad. Sci. 109, 17418–17423.

Schmitt, A., Hamann, F., Neumann, P., and Ficner, R. (2018). Crystal structure of the spliceosomal DEAH-box ATPase Prp2. Acta Crystallogr. Sect. Struct. Biol. 74, 643–654.

Singleton, M.R., Dillingham, M.S., and Wigley, D.B. (2007). Structure and Mechanism of Helicases and Nucleic Acid Translocases. Annu. Rev. Biochem. 76, 23–50.

Sparta, K.M., Krug, M., Heinemann, U., Mueller, U., and Weiss, M.S. (2016). XDSAPP2.0. J. Appl. Crystallogr. 49, 1085–1092.

Tauchert, M.J., Fourmann, J.-B., Lührmann, R., and Ficner, R. (2017). Structural insights into the mechanism of the DEAH-box RNA helicase Prp43. ELife 6, e21510.

Trowitzsch, S., Bieniossek, C., Nie, Y., Garzoni, F., and Berger, I. (2010). New baculovirus expression tools for recombinant protein complex production. J. Struct. Biol. 172, 45–54.

Wan, R., Yan, C., Bai, R., Wang, L., Huang, M., Wong, C.C.L., and Shi, Y. (2016). The 3.8 Å structure of the U4/U6.U5 tri-snRNP: Insights into spliceosome assembly and catalysis. Science 351, 466–475.

Woodman, I.L., Briggs, G.S., and Bolt, E.L. (2007). Archaeal Hel308 domain V couples DNA binding to ATP hydrolysis and positions DNA for unwinding over the helicase ratchet. J. Mol. Biol. 374, 1139–1144.

Xu, D., Nouraini, S., Field, D., Tang, S.J., and Friesen, J.D. (1996). An RNA-dependent ATPase associated with U2/U6 snRNAs in pre-mRNA splicing. Nature 381, 709–713.

Ye, J., Osborne, A.R., Groll, M., and Rapoport, T.A. (2004). RecA-like motor ATPases—lessons from structures. Biochim. Biophys. Acta BBA - Bioenerg. 1659, 1–18.

Zhan, X., Yan, C., Zhang, X., Lei, J., and Shi, Y. (2018). Structures of the human pre-catalytic spliceosome and its precursor spliceosome. Cell Res. 28, 1129–1140.

Zhang, L., Li, X., Hill, R.C., Qiu, Y., Zhang, W., Hansen, K.C., and Zhao, R. (2015). Brr2 plays a role in spliceosomal activation in addition to U4/U6 unwinding. Nucleic Acids Res.

